# Reconstructing Full-Field Flapping Wing Dynamics from Sparse Measurements

**DOI:** 10.1101/2020.06.08.128041

**Authors:** William Johns, Lisa Davis, Mark Jankauski

**Affiliations:** Department of Mathematical Sciences, Montana State University, P.O. Box 172400, Bozeman MT 59717; Department of Mechanical & Industrial Engineering, Montana State University, 220 Roberts Hall, Bozeman MT 59717

## Abstract

Flapping insect wings deform during flight. This deformation benefits the insect’s aerodynamic force production as well as energetic efficiency. However, it is challenging to measure wing displacement field in flying insects. Many points must be tracked over the wing’s surface to resolve its instantaneous shape. To reduce the number of points one is required to track, we propose a physics-based reconstruction method called System Equivalent Reduction Expansion Processes (SEREP) to estimate wing deformation and strain from sparse measurements. Measurement locations are determined using a Weighted Normalized Modal Displacement (NMD) method. We experimentally validate the reconstruction technique by flapping a paper wing from 5-9 Hz with 45° and measuring strain at three locations. Two measurements are used for the reconstruction and the third for validation. Strain reconstructions had a maximal error of 30% in amplitude. We extend this methodology to a more realistic insect wing through numerical simulation. We show that wing displacement can be estimated from sparse displacement or strain measurements, and that additional sensors spatially average measurement noise to improve reconstruction accuracy. This research helps overcome some of the challenges of measuring full-field dynamics in flying insects and provides a framework for strain-based sensing in insect-inspired flapping robots.

## 1. Introduction

The dynamics of flexible insect wings has attracted considerable research interest over the past several decades. During flight, flapping insect wings bend and twist in response to aerodynamic and inertial-elastic forces [1–3]. This deformation is believed to improve the insect’s aerodynamic force production [4] and to enhance its power economy [5–7]. Further, some insects rely on the neurological feedback generated by wing deformation to realize closed-loop attitude control. Hawkmoth *Manduca sexta* wings, for example, are imbued with mechanoreceptors called campaniform sensilla that are believed to trigger reflexive responses when the insect is subject to environmental perturbations [8]. Owing to the substantial role that deformation plays in biological flight, researchers have invested significant effort in measuring wing deformation and flapping kinematics in freely flying insects.

The motion of flapping wings can decomposed into two parts: rigid body motion and elastic deformation. During rigid body motion, all points over the wing’s surface maintain a fixed distance with respect to one another. This occurs during slow articulation of the wing or when the insect’s body translates during flight. During elastic deformation, the relative distance between points is variable, which causes the wing to strain and store elastic potential energy. Given the high frequency flapping of insect flight, both motions occur simultaneously, and the wing’s instantaneous shape and orientation can be determined by superimposing smaller elastic deformation on top of larger rigid body rotation [9, 10]. The concurrence of elastic deformation and rigid body rotation, however, makes it challenging to measure them independently.

The most common mechanism to measure flapping wing motion is through high-speed videography. By utilizing multiple cameras, one can reconstruct the three-dimensional position of a point in space [11]. By tracking multiple points to define time-dependent planes, the kinematics of a structure can be reconstructed. Willmott and Ellington used videography to estimate flapping kinematics of the Hawkmoth *M. sexta* during hovering and forward flight in a wind tunnel [12]. Ortega-Jimenez et al. furthered these studies by measuring *M. sexta* flapping kinematics in turbulence [13]. Greeter and Hedrick later quantified *M. sexta* wing kinematics during lateral maneuvers [14]. Similar kinematic studies have been conducted for honeybees [15], dragonflies [16], as well as other insects [17, 18] and small flapping wing robots inspired by insects [19]. While these studies have undoubtedly advanced our understanding of flapping motions, one potential issue with kinematic reconstruction stems from the motion of the specific points tracked – if these points experience out-of-plane deflection, the resolved kinematics will be influenced by both rigid body rotation and elastic deformation.

Decoupling rigid body rotation from elastic deformation generally requires sophisticated equipment or post-processing techniques in conjunction with high-speed videography. Wu et al. utilized high-speed three-dimensional digital image correlation (DIC) to measure full-field wing deformation in a flapping wing micro air vehicle (FWMAV) in air and in vacuo [20]. Hsu et al. recorded freely flying *M. sexta* and estimated their wing deformation through a complex optimization routine interfaced to a finite element (FE) reference body [21]. Aguayo et al. measured out-of-plane deformation in swallowtail butterfly wings using digital holographic inferometry [22]. Koehler et al. estimated wing deformation in free flying dragonflies using videography and a template-based subdivision surface reconstruction [23]. Unlike kinematic reconstructions, which require only a handful of measurements to define planes, measuring out-of-plane elastic deformation necessitates many measurement points to resolve the wing’s instantaneous shape. Full-field deformation measurements consequently demand significant data storage capabilities, and potentially large amounts of effort to post-process this data. For these reasons, full-field measurements are challenging and often impractical.

To overcome these complications, researchers often estimate full-field quantities based upon sparse measurements using physics-based reconstruction methods. Some of these reconstruction methods include Guyan Condensation, Improve Reduced System, and System Equivalent Reduction Expansion Processes [24]. Each of these methods requires a knowledge of the structure’s mass and stiffness, typically determined either through FE modeling or experimental measurements, but none require explicit knowledge of the structure’s forcing input. This makes them useful when a structure is subject to uncertain loading conditions. Such physics-based reconstruction methods can be employed on flapping wings to reduce the measurement points required to estimate full-field deformation.

### 1.1. Research Scope

The purpose of the present research is to demonstrate that sparse measurements and physics-based reconstruction techniques can be used to estimate full-field flapping wing dynamics. We show this both through a simplified physical experiment and more complex numerical simulation. To the best of our knowledge, such techniques have not previously been applied to flapping wings. Flapping wings present a unique challenge to reconstruction methods because their stiffness varies periodically throughout a wingbeat [9]. From the experimental biology perspective, this research delivers a technique that can reduce the number of points required to track when measuring wing deformation. From the robotics perspective, this research provides a simple framework to estimate instantaneous wing shape on FWMAVs in real-time if strain or displacement is measurable at select points. Instantaneous wing shape, combined with angular rates of rotation, can be used to predict aerodynamic vectors.

The remainder of this paper is organized as follows. First, we summarize System Equivalent Reduction Expansion Processes (SEREP) and Normalized Modal Displacement Method (NMD), the frameworks used to estimate wing dynamics and to optimize sensor placement, respectively. We then introduce an experiment where we flap a paper wing and predict strain at one location using measurements from two others via SEREP. Though we consider only single-degree-of-freedom (SDOF) flapping kinematics and measure strain rather than displacement, our methods our general and scalable to more complex cases. To demonstrate this, we numerically simulate the response of a more realistic insect wing FE model subject to multiple-degree-of-freedom (MDOF) flapping kinematics and reconstruct elastic deformation based on sparse measurements. We conclude by discussing the applicability of this work and the areas of research it can benefit.

## 2. Theory

Here, we introduce the theoretical frameworks utilized throughout this research. Referenced from a coordinate system that rotates with the wing’s rigid body motion (Fig. 1), the spatiotemporal wing deformation *W*(**r**, *t*) is

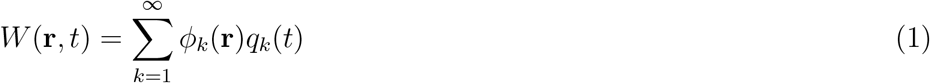

where the vector **r** is the planar coordinate of a differential mass element on the wing, *ϕ*_*k*_ are the wing’s mass normalized mode shapes, and the time dependent modal response is *q*_*k*_(*t*). A similar expansion can be derived for full-field strain replacing the mode shapes *ϕ*_*k*_ with the modal strains. This expansion, which allows the displacement field to be represented in terms of shapes rather than individual points, provides the foundation for the reconstruction methods that follow. Mode shapes *ϕ*_*k*_(**r**) and modal strains can be obtained experimentally or from a finite element discretization of the wing.

**Figure 1:**
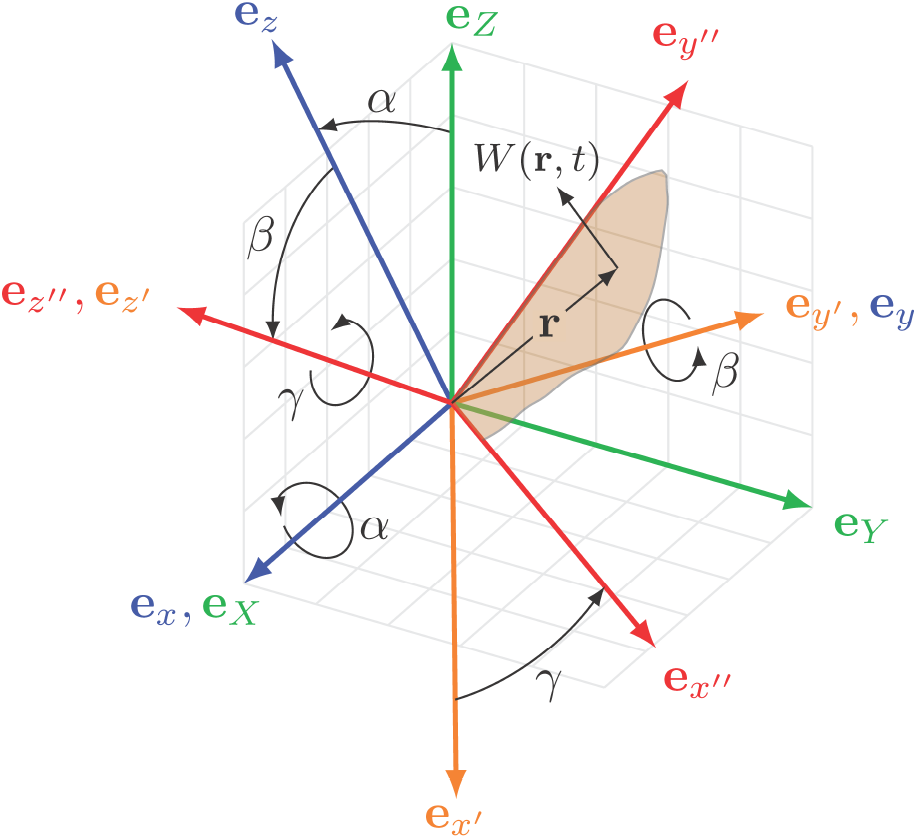
An *X* − *Y* − *Z* inertial reference frame undergoes an *X* − *y* − *z*′ rotation sequence with rotation amplitudes *α*, *β*, *γ*, respectively, where *α* indicates wing roll (flap), *β* indicates wing pitch (rotation), and *γ* indicates wing yaw (stroke deviation or elevation). The terminal *x*′′ − *y*′′ − *z*′′ reference frame is fixed to the rigid body motion of the wing. Out-of-plane elastic deformation *W*(**r**, *t*) occurs in the *z*′′ direction.

### 2.1. System Equivalent Reduction Expansion Process

Model reduction techniques are used to reduce the complexity of systems while retaining important characteristics and for reconstructing systems from sparse measurements. For this work, SEREP is used to reconstruct the full system response from a limited number of measurements [25]. This technique has proven effective for the larger fixed wings on airplanes [26]. We begin by briefly describing the theory underlying SEREP in the current context.

Though continuous in nature, insect wings can practically be treated as MDOF systems if represented by discrete measurement points. We consider reconstructions of **W**, where **W** is the time-dependent vector of out-of-plane elastic deformations referenced from discrete measurement points. Although we show this methodology for displacement reconstructions, strain reconstructions are equally valid. As an MDOF system, wing deformation **W** is

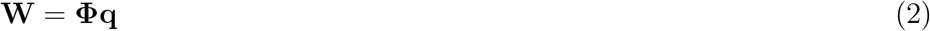

where **Φ** contains the mode shape *ϕ*_*k*_ in the *kth* column and **q** is the modal response vector. We choose a subset of the coordinates to retain, denoted with the subscript *r*, and obtain the reduced system

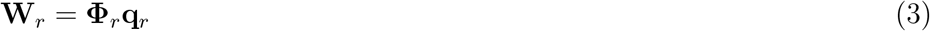

These retained coordinates correspond to the locations of sensors. Information from these sensors is used to reconstruct the full system. For a well defined problem we will need to retain at least as many coordinates as we retain modes in 1. We estimate the full displacement using the generalized inverse of **Φ**_*r*_, denoted by

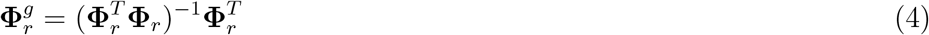

to compute the least squares approximation of W given by

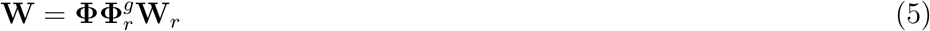

In the special case where we have the same number of retained modes as sensors the generalized inverse is the inverse of **Φ**_*r*_. Error in the measurements **W**_*r*_ may make it beneficial to have more sensors than retained modes.

As modal contributions are shared between strains, it is also possible to mix measurements in different directions to recover the full field strain. For example, ***ϵ***_*x*_ can be reconstructed by

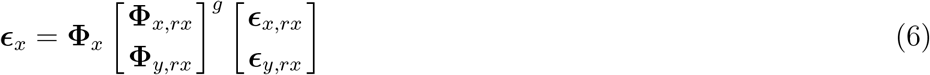

where *rx* and *ry* are the measured coordinates in *x* and *y* respectively and **Φ**_*x*_ and **Φ**_*y*_ contain the modal strains in *x* and *y* respectively. Once the sensor locations are chosen, the mode shapes are used to construct a transformation matrix involving the generalized inverse matrix. This transformation matrix allows us to estimate the full field displacement or strain using 5 and the sensor data. Note that the transformation matrix depends only on the physical parameters of the wing and needs to be computed only once.

### 2.2. Weighted Normalized Modal Displacement Method

Sensor placement can have a large impact on the accuracy of reconstruction. We use the weighted normalized displacement method (NMD) to determine sensor placement [27]. We first compute the driving point residues (DRP) for each node at location **r** over all retained modes *k* by

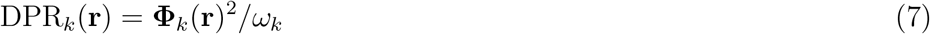

were *ω*_*k*_ is the natural frequency of the *kth* retained mode with mode shape **Φ**_*k*_. We desire sensors with high average DPR (equivalently high modal participation factors) and where each mode contributes significantly. To that end we compute a weighted NMD as

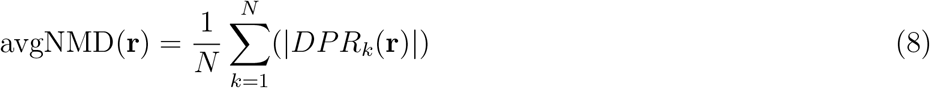

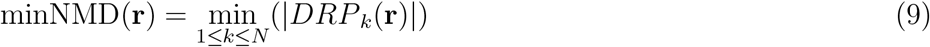

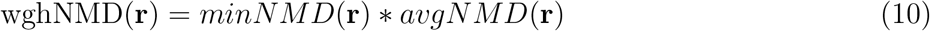

Above, *N* is the total number of measurement points from which *W* is referenced. We select the measurement points with the highest wghNMD as our sensor locations. This formulation can be used to choose strain sensors locations by replacing the mode shapes with the modal strains.

## 3. Experimental Validation

In this section, we present an experiment designed to verify the accuracy of SEREP applied to a simple flapping wing. First, we describe an SDOF mechanism used to flap a wing at 45° between 5-9 Hz. Next, we detail the paper wing used in all experimental trials, where three strain gages are used to measure the wing’s dynamic response. The paper wing is structurally modeled using the finite element method to determine its vibration mode shapes and modal strains. We then estimate wing strain measured at one of the gage locations based on measurements from the other two locations via SEREP.

### 3.1. Mechanical Flapper

The flapping mechanism (Fig. 2) described here originated in [28] and is summarized to provide context to this work. A 60 watt direct current motor (Maxon Motors, 310007) regulated by a proportional-integral-derivative controller (Maxon Motors, EPOS 24/5) drives the wing’s flapping motion. The motor attaches to a fixture that clamps the base of the wing. The end of the clamping fixture is supported by a flange bearing to allow it to spin freely, and the shaft position is measured using a quantized analog encoder (US Digital, MAE3-A10-250-220-7-B). All motor brackets and fixtures are printed using a Form2 SLA 3D printer. We consider flapping frequencies from 5-9 Hz in 1 Hz intervals at an amplitude of 45°. Each flapping trial lasts 15 seconds, where the first 5 seconds are discarded from experimental results to eliminate transient responses. For high-speed video of the flapping mechanism, the reader is directed to the supplementary material. All experimental data is recorded using a National Instrument cDAQ-9178 acquisition system.

**Figure 2:**
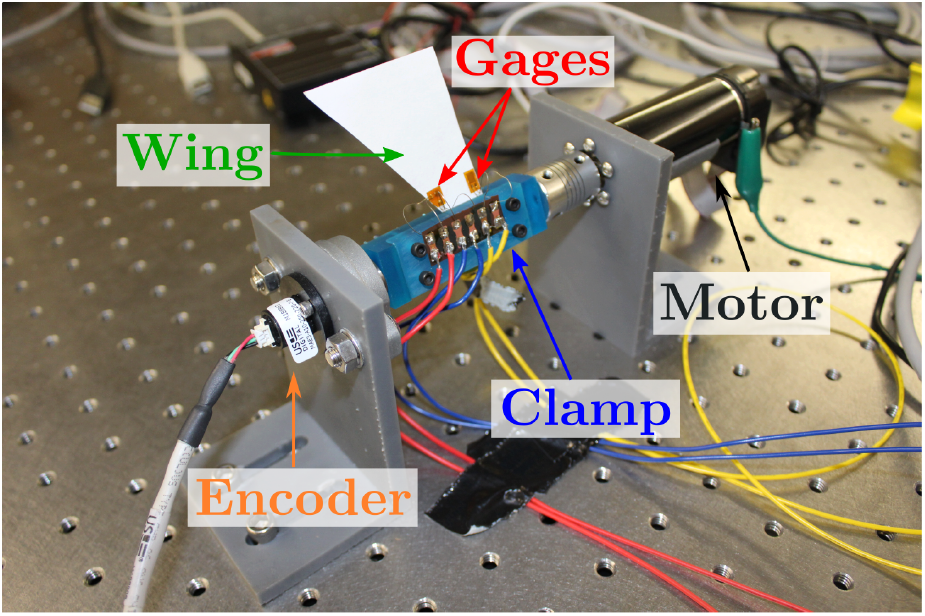
Diagram of flapping mechanism used for experimental trials. The mechanism flaps the paper wing from 5-9 Hz with an amplitude of 45°.

### 3.2. Experimental Wing and Strain Measurements

We create a simple paper wing for experimental trials (Fig. 3). Wing properties are shown in Tab. 1. The wing is made of card stock because it has low surface density and as a result, is influenced by both aerodynamic and inertial loads. The wing is triangular so that it will both bend and twist due to the offset center of mass. Bending will predominately cause the wing to strain in the *Y* direction, whereas twisting (also torsion) causes the wing to strain in the *X* direction. We mount strain gages to the wing using etyhl-based cyanoacrylate, where the strain gage locations are determined by NMD. Due to the physical size of the sensors, one sensor was selected and then moved slightly away from the edge of the wing. The second sensor was then chosen from locations where the sensors would not overlap. We mount one uni-axial gage (Omega Engineering, SGD-2/350-LY11) to measure strain in the *Y* (designated *U*_*y*_), and one bi-axial gage (Omega Engineering, SGD-2/350-XY11) at a separate location that measures strain in both *X* and *Y* directions (designated *B*_*x*_ and *B*_*y*_, respectively). All strain data is recorded using a bridge input module (National Instruments, NI-9236).

**Table 1:** Properties of experimental wing.

| Description | Value | Unit |
| --- | --- | --- |
| Wing Unclamped Length | 5 | cm |
| Wing Width | 5 | cm |
| Wing Thickness | 0.17 | mm |
| Wing Young's Modulus | 9.5 | GPa |
| Wing Poisson Ratio | 0.33 | - |
| Gage Thickness | 0.13 | mm |
| Gage Young's Modulus | 2.5 | GPa |
| Gage Poisson Ratio | 0.33 | - |
| Total Mass | 0.33 | grams |

**Figure 3:**
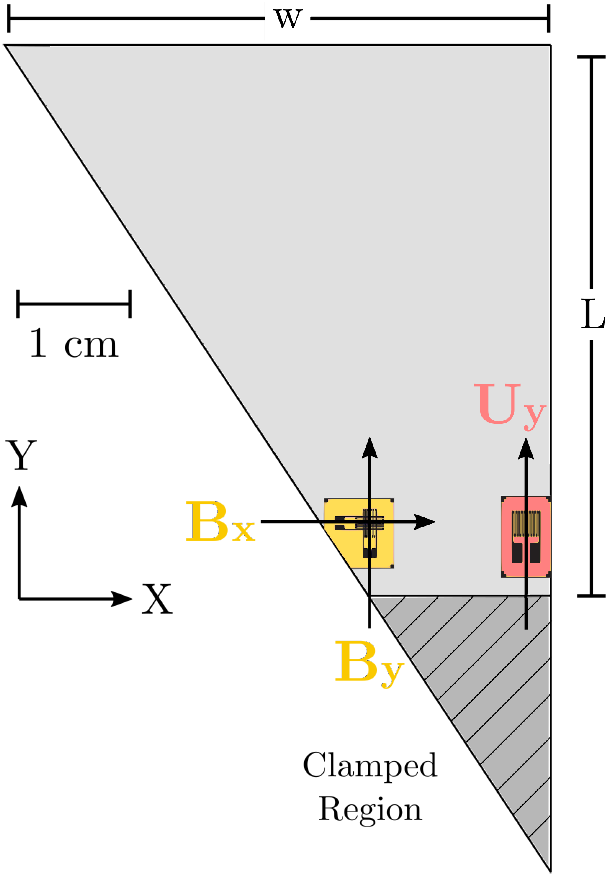
Schematic of the experimental paper wing. Wing is triangular in geometry to promote both twisting and bending during SDOF flapping. *U*_*y*_ designates strain measurement of uni-axial gage in the *Y* direction and *B*_*x*_, *B*_*x*_ designate strain measurements of bi-axial gage in the *X* and *Y* directions respectively. *L* and *w* indicate the wing length and width respectively.

The total of three measurement points implies that we can maximally retain three vibration modes for wing shape reconstruction. However, given that we will utilize only two measurements for reconstruction and the third for validation, we will retain at a maximum two modes in this experimental work. These modes are not required to begin at the first mode, nor do modes need to be adjacent – it is possible to retain the second and fourth mode, for example. Selecting the modes to retain depends on several factors, including wingbeat frequency and the spatiotemporal distribution of aerodynamic forces. For the purposes of this work, we retain either one or two vibration modes in order to demonstrate how this quantity affects the accuracy of the reconstructed signal. We always consider two measurement points. We do not consider any scenarios in which we retain more modes than measurement points, as this represents an underdetermined system of equations. While it is possible to employ SEREP even when the number of modes exceeds number of measurement points by solving a least squares optimization problem, this typically produces undesirable results.

### 3.3. Experimental Wing Structural Modeling

To calculate the vibration modes for SEREP reconstruction, we create an FE model of the wing using ABAQUS. Because the strain gages are of comparable thickness to the paper, they stiffen the overall structure and must be included in modeling efforts. Optimal sensor locations were determined before the gages were included in the FE model; it is possible that inclusion of the sensors in the FE model modestly affects optimal sensor placement. Both the paper wing and gages are modeled using quadrilateral shell elements and the material properties in Tab. 1. The wing is discretized into 884 elements total, which is sufficient for convergence of the wing’s first four natural frequencies. We then conduct a modal analysis to determine the wing’s mode shapes, which includes both information regarding modal displacements and directional modal strains. The first four vibration modes are shown in Fig. 4. From first to fourth, the wing’s natural frequencies are 21.3 Hz, 63.9 Hz, 161.3 Hz and 263.9 Hz respectively. Natural frequencies are not required for SEREP, however they do inform which vibration modes should be retained if the input bandwidth is known. Further, natural frequencies are required to compute DRP (Eq. 9). While we have utilized the finite element method to determine the modal displacements and strains, these quantities may also be identified experimentally if desired.

**Figure 4:**
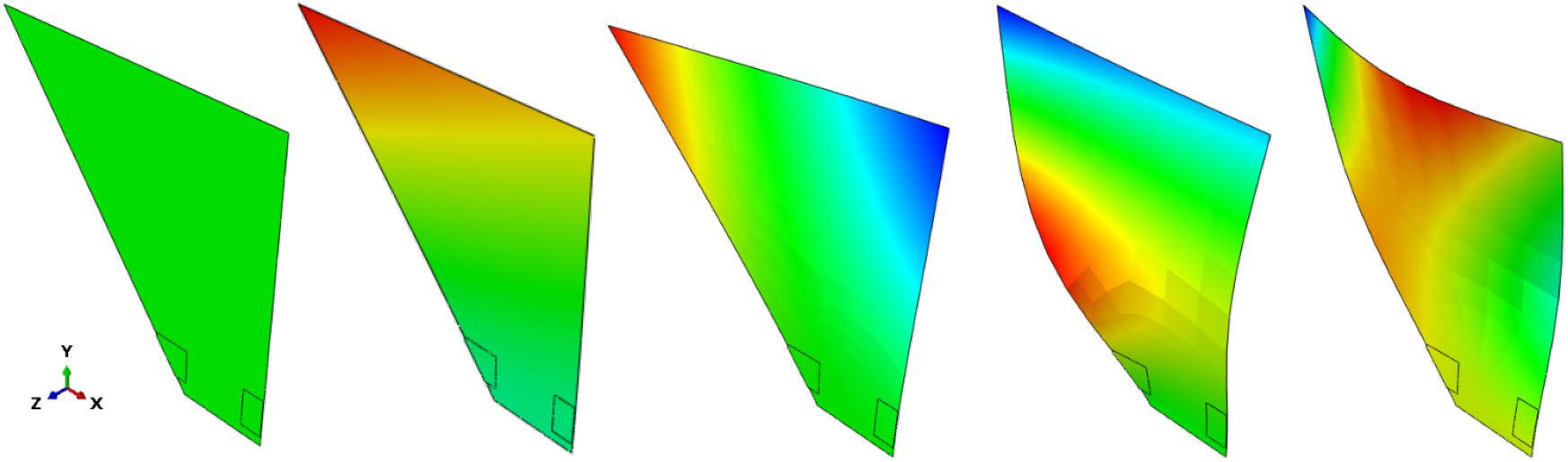
From left to right, finite element model of the experimental wing in its undeformed state followed by the first four vibration modes in ascending order. Color maps indicate relative modal displacement for each vibration mode. Note that modal displacement is a dimensionless quantity, and should only be compared within a single mode rather than across distinct modes.

### 3.4. Experimental Results

Here, we present the results of the SDOF flapping experiments. In general, SEREP is effective at predicting strain from sparse measurements (Figs. 5-6). Prediction accuracy appears independent of excitation frequency, and the method was effective at accounting for qualitative differences between strain waveforms. Retaining more measurement points improved the agreement between the measured and predicted strain. We show only results for 6 and 9 Hz flapping frequencies owing to their dissimilar strain responses; trends in reconstructed strain are similar for intermediate flapping frequencies.

**Figure 5:**
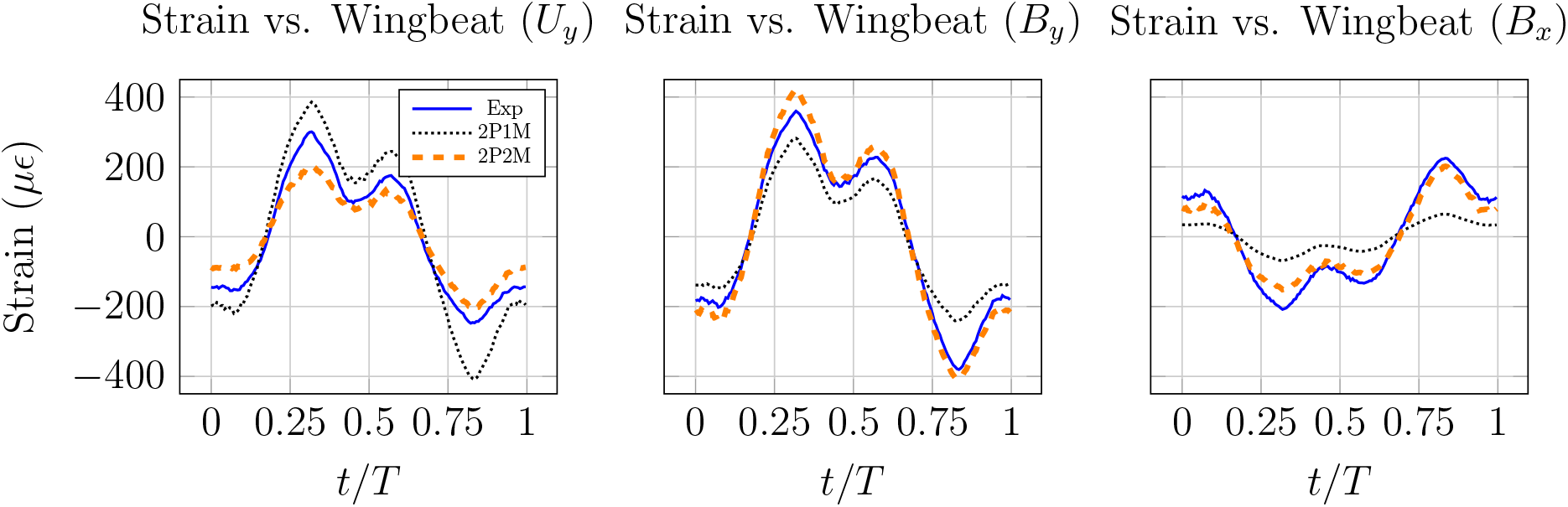
Measured and reconstructed strain for a wing flapping at 6 Hz shown over a single wingbeat. 2P1M indicates a reconstruction with two measurement points, one retained mode, and 2P2M indicates a reconstruction with two measurement points, two retained modes. Note the significant strain response at three times the flapping frequency, which occurs due to proximity to a superharmonic resonance.

**Figure 6:**
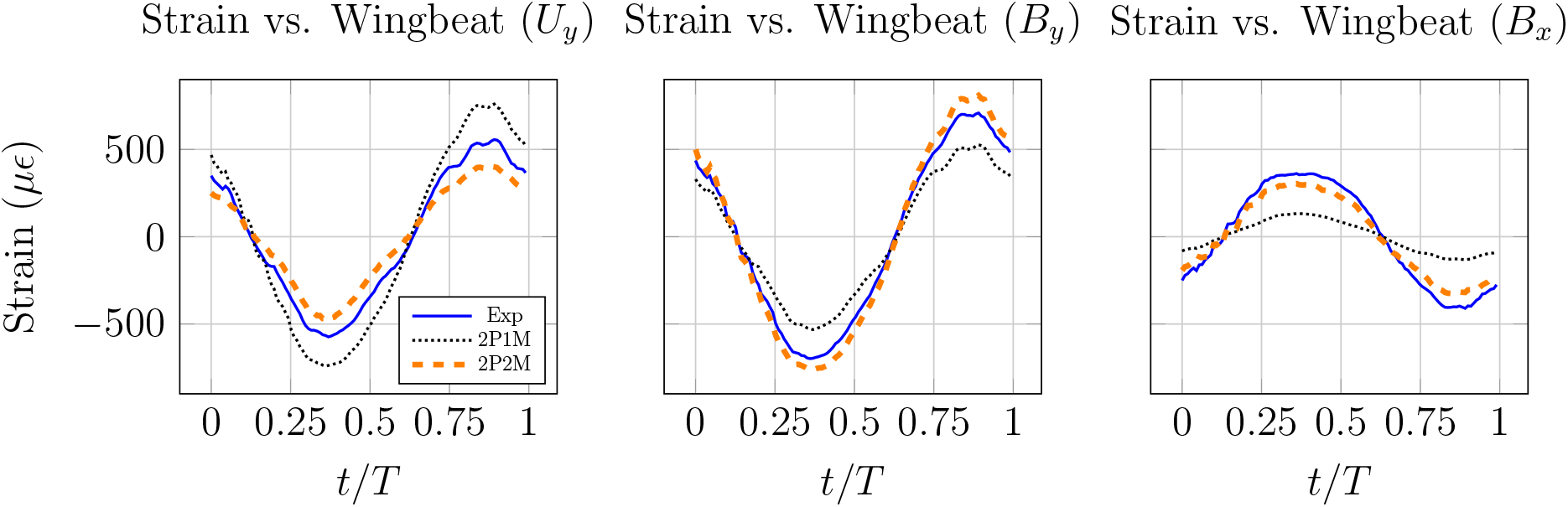
Measured and reconstructed strain for a wing flapping at 9 Hz shown over a single wingbeat. 2P1M indicates a reconstruction with two measurement points, one retained mode, and 2P2M indicates a reconstruction with two measurement points, two retained modes. Compared to the 6 Hz case, the strain response at three times the flapping frequency has diminished significantly.

Interestingly, retention of the first and third vibration modes (Fig. 4) produced the best reconstructed strain signals in all cases. This is likely due to how the aerodynamic force is distributed over the flapping wing. The second vibration mode is a torsional mode, whereas the third vibration mode is a bending mode. The aerodynamic force acting on an SDOF flapping wing will approximately scale quadratically along the *Y* - axis from the axis of rotation (Fig. 3), as well as with the square of angular velocity [28]. Because the wing’s torsional mode exhibits displacement in both the positive and negative *Z* direction, the projection of the aerodynamic force onto the torsional mode will be close to zero. On the other hand, projecting this aerodynamic force onto the first and third vibration modes will generate a non-zero modal excitation. Thus, we believe the wing’s displacement is best described via the first and third vibration modes, despite intentionally offsetting the center of mass to promote a torsional response. For the following results where only one mode is retained, we retain the first vibration mode.

Having identified which modes are most appropriate to retain for SEREP, we turn to the reconstructed strain signals. First, consider the strain profiles depicted in Fig. 5 for a wing flapping at 6 Hz. For all strain sensors, the primary response occurs at the flapping frequency, where *B*_*y*_ and *U*_*y*_ are in-phase and *B*_*x*_ is 180° out-of-phase. Because the wing is flapping at roughly 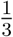 its natural frequency, it experiences a near super-harmonic resonance which contributes to a large strain response at three times the flapping frequency [28]. SEREP captures this effect despite having no direct measurement of the input. The reconstructed strain signals are generally good both in terms of magnitude and phase. Intuitively, the reconstruction is more accurate when more vibration modes are retained. When considering two retained modes, the largest discrepancy between measured and reconstructed strain occurs in *U*_*y*_, which maximally has an error of about 30% in magnitude.

Next, we turn our attention to 9 Hz flapping (Fig. 6). Similar to the 6 Hz case, *B*_*y*_ and *U*_*y*_ are in-phase and *B*_*x*_ is 180° out-of-phase. The strain response at three times the flapping frequency has diminished significantly, because the third harmonic of the flapping frequency is no longer in proximity to the wing’s first resonant frequency. Strain reconstructions are of similar accuracy to the 6 Hz case. The largest error remains in *U*_*y*_, but is reduced by about 5%. The reduction in error likely comes from a change in modal participation factors; it is possible that unretained modes more notably influenced the wing while it was flapping at 6 Hz, but that those modes are not as influential when the flapping frequency is increased to 9 Hz. While inaccuracies in the FE model may cause errors in strain reconstructions, such errors would be systematic and affect the reconstructions at both flapping frequencies equivalently.

## 4. Numerical Simulation

In the previous section, we demonstrated that SEREP effectively reconstructs strain for a simplified flapping wing. However, insect wings represent a more significant challenge. They are made up of a thin membrane reinforced by thicker veins [29], have a cambered surface to promote geometric stiffening [30], and experience MDOF rotational flapping [12]. Further, because the natural frequencies in some insect wings are closely spaced [31], it is likely that several vibration modes contribute to the wing’s overall deformation.

To further investigate SEREP’s ability to estimate full-field flapping wing dynamics, we numerically simulate the response of a more realistic insect wing with conventional flapping kinematics. First, we derive an FE model of an *M. sexta* forewing that considers both camber and venation. Then, we utilize an inertial-elastic model to simulate the full-field response of the flapping wing. Lastly, we examine how sensor type, number and placement influence the accuracy of reconstruction methods.

### 4.1. Simulation Wing and Kinematics

We develop the *M. sexta* forewing model using ABAQUS (Fig. 7). The planar wing geometry is based off a micro computed tomography image from [32]. The membrane is modeled using shell elements and the veins are modeled using beam elements, where vein cross sections are assumed solid and circular and have effective diameters ranging from 100-400 *μ*m. We apply a curvature to both the membrane and veins using reported results from [33]. The membrane and veins are restricted to move together by tie constraints. All material properties for the wing are approximated from [30] and are detailed in Tab. 2. We assume the materials to be linear and isotropic. All degrees of freedom are fixed at the wing root. The model consists of 774 elements in total, which is sufficient for convergence of the wing’s first four natural frequencies.

**Figure 7:**
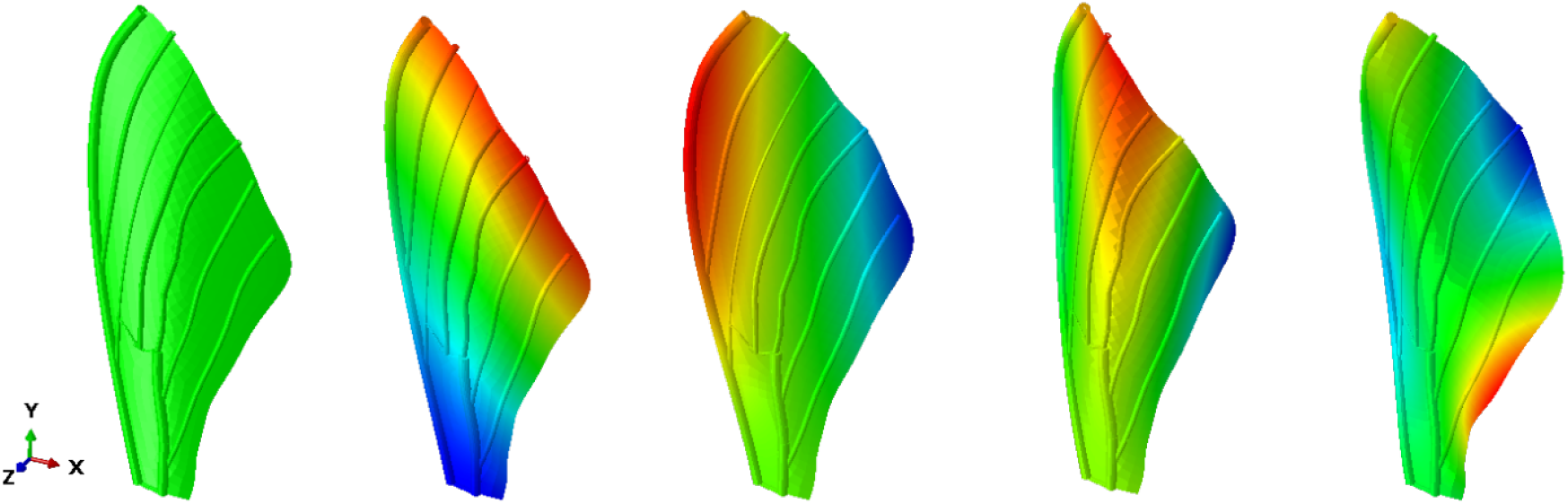
From left to right, finite element model of the idealized insect wing in its undeformed state followed by the first four vibration modes in ascending order. Color maps indicate relative modal displacement for each vibration mode. Vein diameters are magnified by 3x to facilitate visualization.

**Table 2:** Properties of simulation wing.

| Description | Value | Unit |
| --- | --- | --- |
| Membrane Density | 1500 | kg/m<sup>3</sup> |
| Membrane Young’s Modulus | 4.0 | GPa |
| Membrane Poisson Ratio | 0.33 | - |
| Membrane Thickness | 50 | μm |
| Vein Density | 1500 | kg/m<sup>3</sup> |
| Vein Young’s Modulus | 8.0 | GPa |
| Wing Span | 50.5 | mm |
| Wing Max Chord Width | 20 | mm |

With the model built, we conduct a modal analysis to determine the wing’s first four natural frequencies and mode shapes (Fig. 7). The first and second natural frequencies agree fairly well with those reported for *M. sexta* [31], however the third and fourth are considerably off. This is likely due to some of the modeling simplifications, for example linear isotropic material behaviors, as well as the inherent complexity of modeling higher-order vibration modes accurately. We manually adjust the FE wing natural frequencies to agree with reported values prior to calculating the wing displacement during flapping. The original natural frequency values, as well as the adjusted values, are shown in Tab. 3. Lowering the third and fourth natural frequencies to be in closer proximity to the flapping frequency will increase their contributions to the overall wing deformation.

**Table 3:** Natural frequencies of the FE insect wing are calculated and are subsequently adjusted to match measurements reported in the literature [31]. All frequencies are in Hz.

| | $\omega_1$ | $\omega_2$ | $\omega_3$ | $\omega_4$ |
| --- | --- | --- | --- | --- |
| Original Value | 57.9 | 94.4 | 275.4 | 391.5 |
| Adjusted Value | 60 | 84 | 107 | 142 |

Lastly, to estimate wing deformation, we implement the model developed in [34]. This model assumes the rigid body rotations are known at the base of the wing (Fig. 1). We simplify these rotations from [12], and assume a roll amplitude of 60°, a pitch amplitude of 45°, and a pitch-roll phase difference of 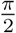 rad. We assume all rotations are harmonic with a flapping frequency of 25 Hz. Yaw rotation angle, also called stroke deviation or elevation, are not considered. To reduce computational costs, we neglect aerodynamic loading. While this will affect the wing’s response, it will not in general affect the accuracy of SEREP or NMD since these methods rely only on the FE model and not dynamic inputs. It is possible that aerodynamics will influence modes not excited by inertial elastic forcing, however the general trends in reconstruction accuracy presented hereafter are expected to be similar.

### 4.2. Simulation Results

We estimate full-field wing displacement using sparse strain measurements and displacement measurements. Sensor locations determined by NMD for both cases are shown in Fig. 8. We place up to sixteen sensors, where sensor one corresponds to the highest priority and sensor sixteen the lowest. For simplicity, we enforce that the sensor placements occur on the vein structure. Strain sensors are assumed to measure bending strain in the vein’s axial direction. We select two points on the wing’s leading and trailing edges to assess reconstruction accuracy, where both points are referenced from the wing’s membrane. Because noiseless measurements result in perfect reconstruction, we add random Gaussian noise of ± 0.5 mm to displacement measurements (≈ 10% maximal displacement) and ± 5 *μϵ* to strain measurements (≈ 0.5% maximal strain) in order to demonstrate the benefit to including additional measurement points in reconstruction efforts. While these noise levels are relatively low, all results in this section are shown unaltered by signal processing, both the measurements used for reconstructions or the reconstructions themselves.

**Figure 8:**
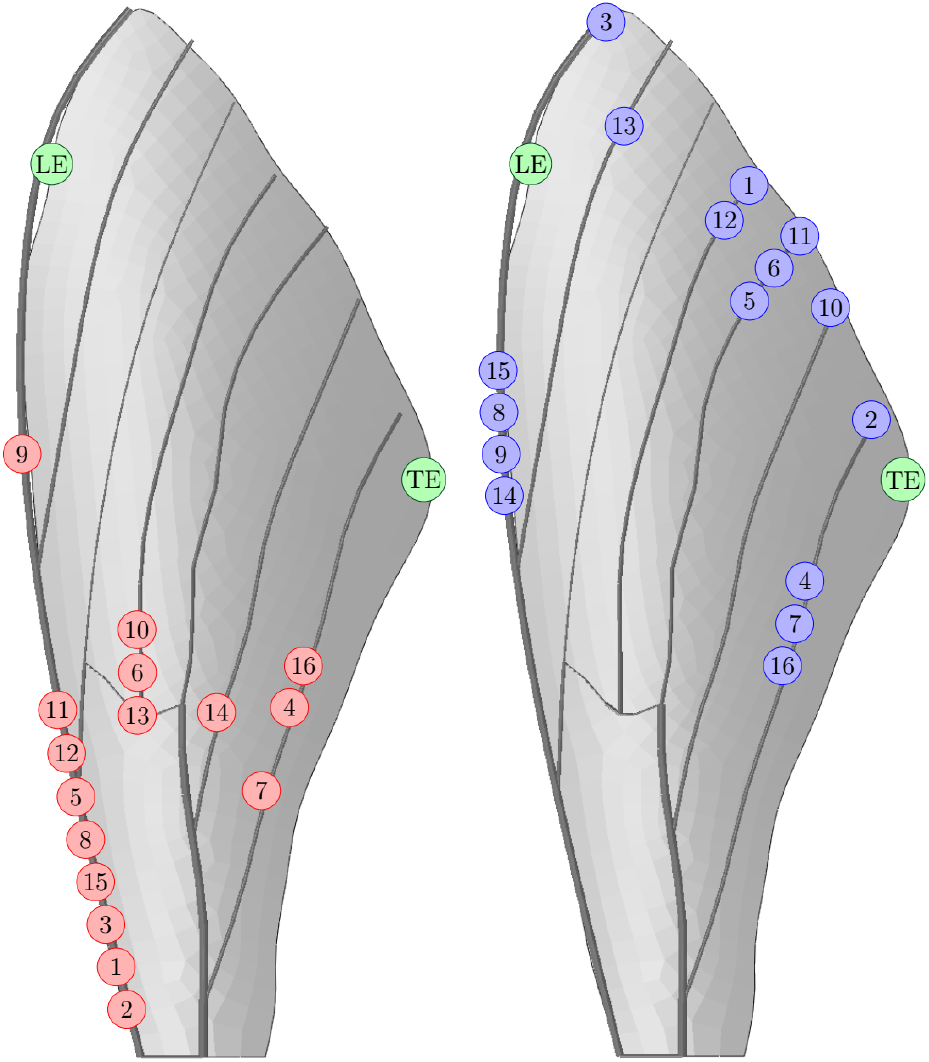
Sensor locations determined by NMD for SEREP reconstruction, where 1 is the first sensor placement and 16 is the last. (Left) Strain measurement locations for strain to displacement reconstruction, and (Right) Displacement measurement locations for displacement to displacement reconstruction.

First, consider the strain sensor placements (Fig. 8, left). The highest priority gages congregate near the base of the leading edge vein where the strain is greatest during flapping. The fourth, sixth and seventh gages are placed near the trailing edge and wing’s torsional lines, which are locations that experience large strains when the wing twists. The remaining gages are placed in similar high strain regions. Next, consider the displacement sensor placements (Fig. 8, right). Most sensors are located in the distal region of the wing, where it experiences the largest deformations. Sensors one through three are located between the wingtip and trailing edge, near the trailing edge and at the wingtip, respectively. Each of these points experience significant motion as the wing’s response tends to be dominated by the first and second vibration modes.

Beyond, sensors tend to cluster in the distal region of the wing, with another grouping emerging on the leading edge vein. While it is possible to define a minimal distance that sensors must be placed with respect to one another to prevent clustering, we did not consider this here.

With measurement points selected, we turn our attention to reconstructed displacements. The leading and trailing edge point reconstructions, superimposed on exact displacements, are shown in Figs. 9-10 for four and sixteen sensors. To quantify the difference between the reconstructed and exact displacements, we define an error proxy as

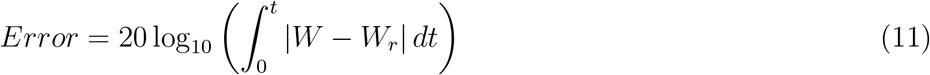

where *W* is evaluated at either the leading or trailing edge and the error units are in decibels. The error is shown as a function of sensor count in Figs. 9-10. For strain reconstructions, SEREP predicts leading edge displacement well for either four and sixteen sensors. This likely has to do with the high concentration of sensors along the leading edge vein, and the proximity of the tracked point to this vein. In contrast, the trailing edge is well-predicted by sixteen sensors but experiences significant noise for four sensors. In this case, many sensors are required to average the measurement noise artificially to produce a good reconstruction, at least without applying a low-pass or moving mean filter to the reconstructed signal. For the displacement reconstructions, the leading edge point requires more sensors to reconcile noise. Again, this is likely because the four initial sensors are relatively distant from the reconstruction point. Trailing edge displacement is reasonably well represented by four sensors, though inclusion of additional sensors does not significantly reduce the noise that does exist. In all cases, increasing sensors decreases error monotonically. It appears that the error at both measurement points will eventually converge asymptotically, though at a greater sensor amount than is considered here.

**Figure 9:**
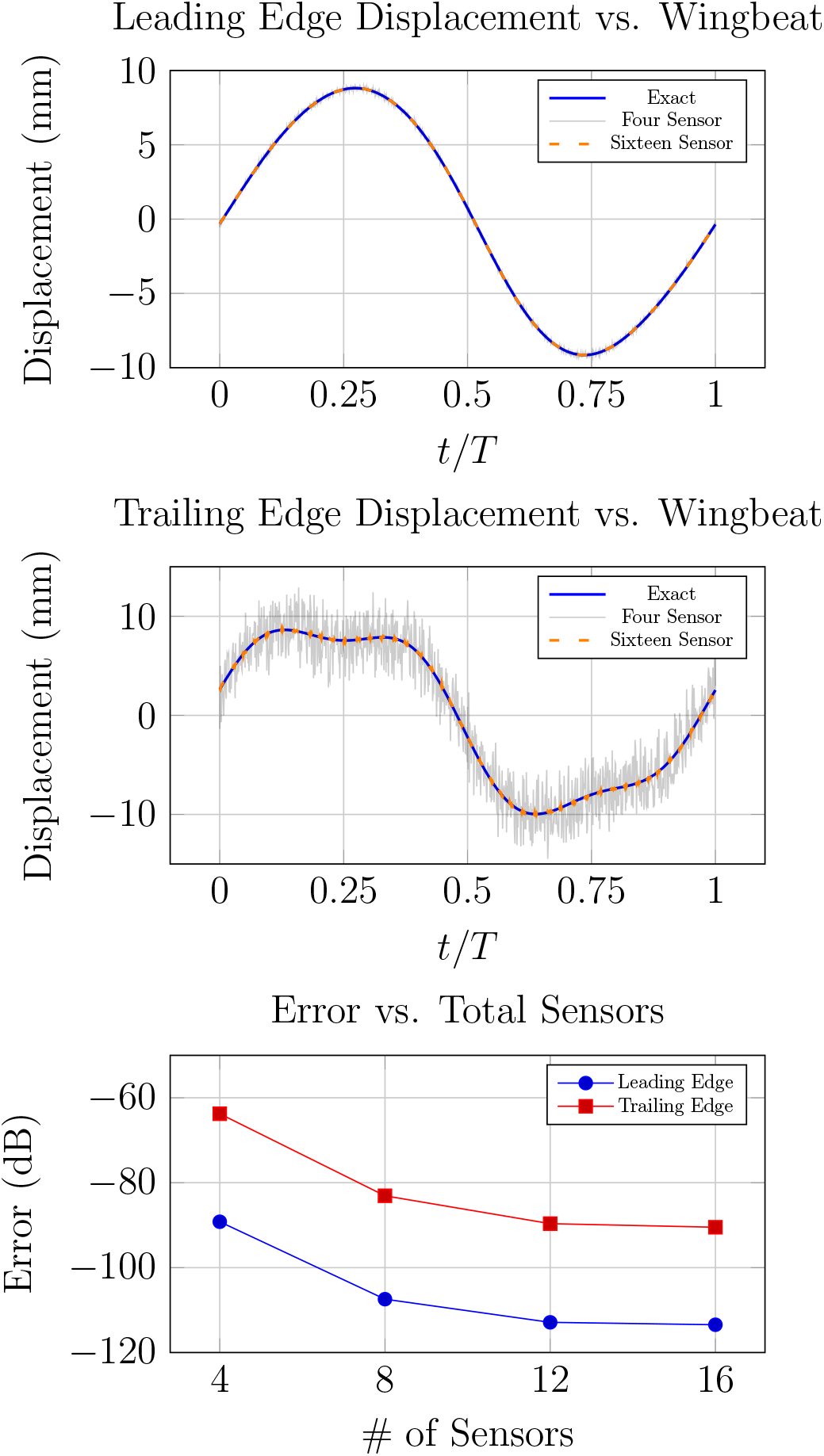
Full-field displacement reconstructions from strain measurements for the simulated insect wing. Top and middle show leading edge and trailing edge displacements and displacement reconstructions for four and sixteen sensors. Bottom shows the error in reconstruction as a function of the number of sensors.

**Figure 10:**
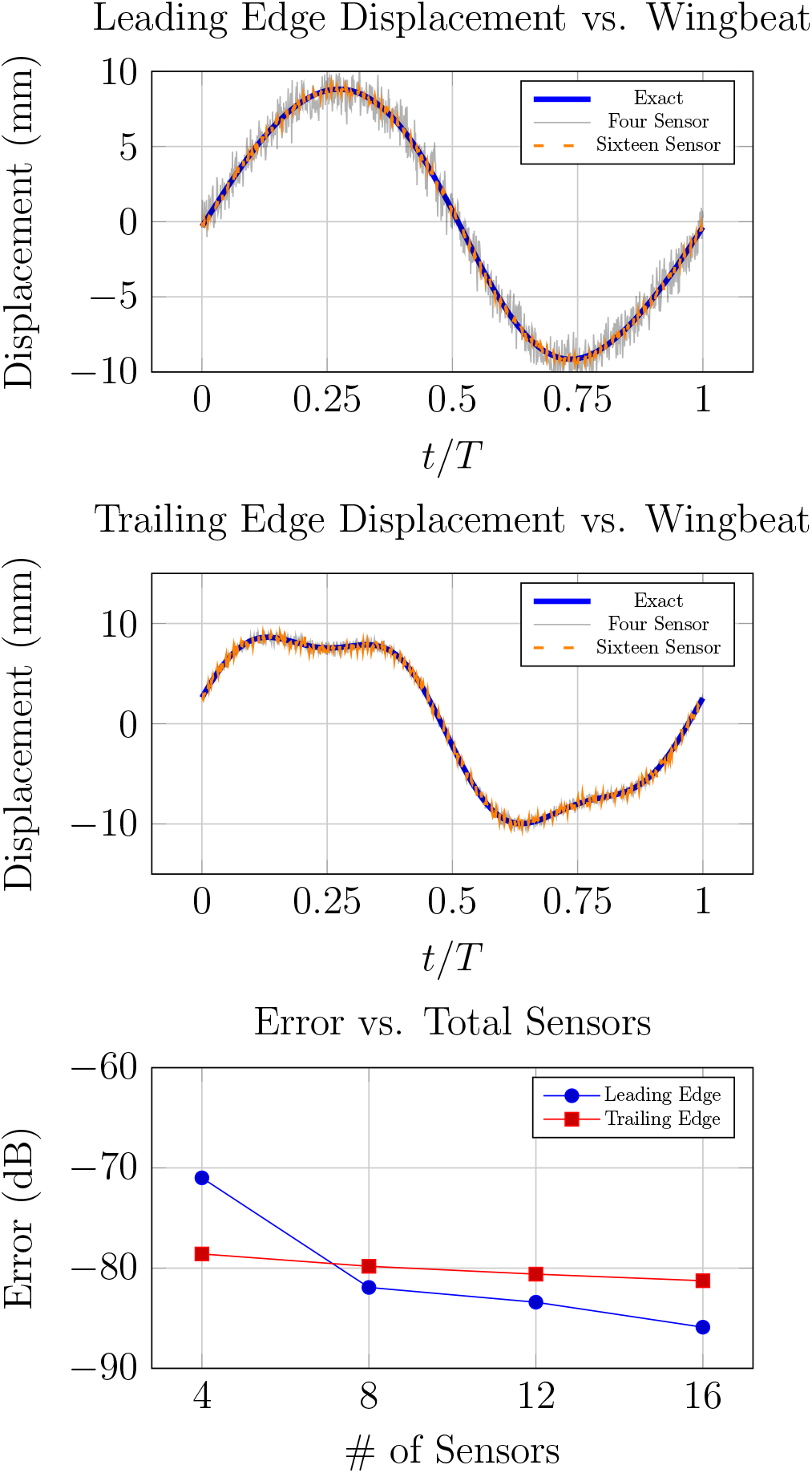
Full-field displacement reconstructions from displacement measurements for the simulated insect wing. Top and middle show leading edge and trailing edge displacements and displacement reconstructions for four and sixteen sensors. Bottom shows the error in reconstruction as a function of the number of sensors.

## 5. Discussion

Deformation is integral to the function of flapping wings, however measuring deformation over an entire wing surface is challenging or impractical in many contexts. Through this research, we demonstrate how SEREP can estimate full-field quantities using sparse measurements, where the optimal locations of the measurement points are informed by NMD. SEREP is general in the sense that it can estimate displacement from strain or visa versa, as well as strain from strain. Through experiment, we show that we are able to estimate a strain waveform at one sensor location based upon two other measurements with good accuracy. Further numerical simulation on a more complex insect wing subject to realistic flapping kinematics showed that wing deformation can effectively be reconstructed from sparse strain or displacement measurements, and that additional sensors reduce reconstruction noise via spatial averaging.

### 5.1. Applications in Biology

The methods developed in this research can help overcome some of the challenges of measuring flapping wing deformation. First, SEREP can help reconcile issues associated with landmark occlusion. Throughout a flight sequence, it is possible that a tracked point moves out of the camera field of view because it is occluded by the insect’s body or other wing. Occlusion can be eliminated if a sufficient number of cameras are used for point tracking, but there tends to be a practical limit due to monetary or spatial constraints. Because SEREP requires sampling at only a handful of points to determine full-field wing deformation, it provides a good mechanism to estimate the wing shape in video frames where measurement points cannot be seen. Second, if an accurate FE model is available, SEREP can facilitate reconstructions inclusive of mesoscale wing features such as wing corrugation and venation. These features are difficult to track without a high density of measurement points [18]. Optical methods have also been proposed as a means to incorporate mesoscale features into wing reconstructions [23]. Lastly, reconstructions based upon sparse measurements reduce the computational time associated with estimating full-field dynamics. SEREP is algebraic, and the reconstruction matrix in Eq. 5 is pre-computed from the wing’s stationary vibration modes. As a result, reconstructions can be conducted efficiently, including in real time if desired.

Nonetheless, there are limitations to the methods presented in this research as well. Most notably, both SEREP and NMD require the wing’s vibration modes to be known. Modes are typically approximated through FE modeling or measured directly via experimental modal analysis. While wing mode shapes and FE models are available for the most commonly studied flying insects such as the *M. sexta* [30, 31], they are not readily accessible for less common insects. FE models are challenging to develop for some insects due to a lack of information regarding geometric and material properties. Measuring vibration modes experimentally is straightforward, however these modes are not scaled with respect to the wing’s mass. Mass scaling is not required by SEREP, though many sensor placement algorithms necessitate vibration modes to be normalized with respect to the wing’s mass. While there are experimental techniques to estimate mass normalized modes, they often require systematic addition and removal of weight to the structure [35] which may be prohibitive for lightweight wings.

### 5.2. Applications in Robotics

The instantaneous wing shape of small FWMAVs influences the vehicle’s aerodynamics. If strain is measurable, SEREP can be used to estimate this shape. Strain gages themselves are infeasible for centimeter-scale aircraft – in addition to adding weight and affecting passive wing deformation, the wiring network would quickly become cumbersome if multiple sensors were employed. On the other hand, advances in printable piezoelectrics may soon enable both the strain sensor and wiring network to be printed directly to the wing [36]. This would facilitate inclusion of dense sensor networks, similar to those observed in real insect wings [8]. In addition to wing shape, these networks of strain sensors may also facilitate identification of aerodynamic environment the vehicle is navigating [37].

Of course, instantaneous shape alone is not sufficient to predict aerodynamic forces. This would require additional knowledge of the flapping kinematics, such as the components of angular velocity. While we have not attempted to predict rotation rates from strain in this work, Eberle et al. showed that this mapping is possible [38]. However, even if wing shape and rotation rates are known, flapping wing fluid-structure interaction (FSI) models are still too high-order to make predictions in real time; most models capable of predicting aerodynamics efficiently neglect wing deformation altogether [39–41]. Though flapping wing FSI models continue to improve, most typically they employ limiting assumptions such as a rigid leading edge [42] or uni-lateral coupling between fluid and structure [34]. Nonetheless, the research presented here provides an approach that may ultimately improve aerodynamic sensing in FWMAVs moving forward.

## References

[1] Bruno A Roccia, Sergio Preidikman, and Balakumar Balachandran. Computational dynamics of flapping wings in hover flight: a co-simulation strategy. AIAA Journal, 55(6):1806–1822, 2017.

[2] T Fitzgerald, M Valdez, M Vanella, E Balaras, and B Balachandran. Flexible flapping systems: computational investigations into fluid-structure interactions. The Aeronautical Journal, 115(1172):593–604, 2011.

[3] Kenji Takizawa, Tayfun E Tezduyar, and Nikolay Kostov. Sequentially-coupled space–time fsi analysis of bio-inspired flapping-wing aerodynamics of an mav. Computational Mechanics, 54(2):213–233, 2014.

[4] Andrew M Mountcastle and Stacey A Combes. Wing flexibility enhances load-lifting capacity in bumblebees. Proceedings of the Royal Society B: Biological Sciences, 280(1759):20130531, 2013.

[5] Bo Yin and Haoxiang Luo. Effect of wing inertia on hovering performance of flexible flapping wings. Physics of Fluids, 22(11):111902, 2010.

[6] Heidi E Reid, Ryan K Schwab, Miles Maxcer, Robert KD Peterson, Erick L Johnson, and Mark Jankauski. Wing flexibility reduces the energetic requirements of insect flight. Bioinspiration & biomimetics, 14(5):056007, 2019.

[7] Mark Jankauski, Ziwen Guo, and IY Shen. The effect of structural deformation on flapping wing energetics. Journal of Sound and Vibration, 429:176–192, 2018.

[8] Bradley H Dickerson, Zane N Aldworth, and Thomas L Daniel. Control of moth flight posture is mediated by wing mechanosensory feedback. Journal of Experimental Biology, 217(13):2301–2308, 2014.

[9] Mark Jankauski and IY Shen. Dynamic modeling of an insect wing subject to three-dimensional rotation. International Journal of Micro Air Vehicles, 6(4):231–251, 2014.

[10] Mark Jankauski and IY Shen. Experimental studies of an inertial-elastic rotating wing in air and vacuum. International Journal of Micro Air Vehicles, 8(2):53–63, 2016.

[11] Tyson L Hedrick. Software techniques for two-and three-dimensional kinematic measurements of biological and biomimetic systems. Bioinspiration & biomimetics, 3(3):034001, 2008.

[12] Alexander P Willmott and Charles P Ellington. The mechanics of flight in the hawkmoth manduca sexta. i. kinematics of hovering and forward flight. Journal of experimental Biology, 200(21):2705–2722, 1997.

[13] Victor Manuel Ortega-Jimenez, Jeremy SM Greeter, Rajat Mittal, and Tyson L Hedrick. Hawkmoth flight stability in turbulent vortex streets. Journal of Experimental Biology, 216(24):4567–4579, 2013.

[14] Jeremy SM Greeter and Tyson L Hedrick. Direct lateral maneuvers in hawkmoths. Biology open, 5(1):72–82, 2016.

[15] Douglas L Altshuler, William B Dickson, Jason T Vance, Stephen P Roberts, and Michael H Dickinson. Short-amplitude high-frequency wing strokes determine the aerodynamics of honeybee flight. Proceedings of the National Academy of Sciences, 102(50):18213–18218, 2005.

[16] Hao Wang, Lijiang Zeng, Hao Liu, and Chunyong Yin. Measuring wing kinematics, flight trajectory and body attitude during forward flight and turning maneuvers in dragonflies. Journal of Experimental Biology, 206(4):745–757, 2003.

[17] Shih-Jung Hsu, Neel Thakur, and Bo Cheng. Speed control and force-vectoring of bluebottle flies in a magnetically levitated flight mill. Journal of Experimental Biology, 222(4):jeb187211, 2019.

[18] Simon M Walker, Adrian LR Thomas, and Graham K Taylor. Photogrammetric reconstruction of high-resolution surface topographies and deformable wing kinematics of tethered locusts and free-flying hoverflies. Journal of the royal society interface, 6(33):351–366, 2009.

[19] John P Whitney and Robert J Wood. Aeromechanics of passive rotation in flapping flight. Journal of Fluid Mechanics, 660:197–220, 2010.

[20] P Wu, B Stanford, W Bowman, A Schwartz, and P Ifju. Digital image correlation techniques for full-field displacement measurements of micro air vehicle flapping wings. Experimental Techniques, 33(6):53–58, 2009.

[21] Tzu-Sheng Shane Hsu, Timothy Fitzgerald, Vincent Phuc Nguyen, Trisha Patel, and Balakumar Balachandran. Motion visualization and estimation for flapping wing systems. Acta Mechanica Sinica, 33(2):327–340, 2017.

[22] Daniel D Aguayo, Fernando Mendoza Santoyo, H Manuel, Manuel D Salas-Araiza, Cristian Caloca-Mendez, and David Asael Gutierrez Hernandez. Insect wing deformation measurements using high speed digital holographic interferometry. Optics express, 18(6):5661–5667, 2010.

[23] Christopher Koehler, Zongxian Liang, Zachary Gaston, Hui Wan, and Haibo Dong. 3d reconstruction and analysis of wing deformation in free-flying dragonflies. Journal of Experimental Biology, 215(17):3018–3027, 2012.

[24] Peter Avitabile. Model reduction and model expansion and their applications–part 1 theory. In Proceedings of the Twenty-Third International Modal Analysis Conference, Orlando, FL, USA, 2005.

[25] J O’Callahan, P Avitabile, and R Riemer. System equivalent reduction expansion process. 01 1988.

[26] Chan-Gi Pak. Wing shape sensing from measured strain. 01 2015.

[27] Daniel C. Kammer. Sensor placement for on-orbit modal identification and correlation of large space structures. Journal of Guidance, Control, and Dynamics, 14(2):251–259, 1991.

[28] Ryan K Schwab, Heidi E Reid, and Mark Jankauski. Reduced-order modeling and experimental studies of bilaterally coupled fluid–structure interaction in single-degree-of-freedom flapping wings. Journal of Vibration and Acoustics, 142(2), 2020.

[29] Y Meresman, JF Husak, R Ben-Shlomo, and G Ribak. Morphological diversification has led to inter-specific variation in elastic wing deformation during flight in scarab beetles. Royal Society Open Science, 7(4):200277, 2020.

[30] SA Combes and TL Daniel. Flexural stiffness in insect wings i. scaling and the influence of wing venation. Journal of experimental biology, 206(17):2979–2987, 2003.

[31] Aaron G Norris, Anthony N Palazotto, and Richard G Cobb. Experimental structural dynamic characterization of the hawkmoth (manduca sexta) forewing. International Journal of Micro Air Vehicles, 5(1):39–54, 2013.

[32] Aaron G Norris. Experimental characterization of the structural dynamics and aero-structural sensitivity of a hawkmoth wing toward the development of design rules for flapping-wing micro air vehicles. Air Force Institute of Technology, 2013.

[33] RP O’Hara and AN Palazotto. The morphological characterization of the forewing of the manduca sexta species for the application of biomimetic flapping wing micro air vehicles. Bioinspiration & biomimetics, 7(4):046011, 2012.

[34] Ryan Schwab, Erick Johnson, and Mark Jankauski. A novel fluid–structure interaction framework for flapping, flexible wings. Journal of Vibration and Acoustics, 141(6), 2019.

[35] P Fernández, Paul Reynolds, and Manuel López-Aenlle. Scaling mode shapes in output-only systems by a consecutive mass change method. Experimental mechanics, 51(6):995–1005, 2011.

[36] Weiwei Xu, Hsien-Lin Huang, Yifeng Liu, Chuan Luo, GZ Cao, and IY Shen. Fabrication and characterization of pzt-silane nano-composite thin-film sensors. Sensors and Actuators A: Physical, 246:102–113, 2016.

[37] Krithika Manohar, Steven L Brunton, and J Nathan Kutz. Environment identification in flight using sparse approximation of wing strain. Journal of Fluids and Structures, 70:162–180, 2017.

[38] AL Eberle, BH Dickerson, PG Reinhall, and TL Daniel. A new twist on gyroscopic sensing: body rotations lead to torsion in flapping, flexing insect wings. Journal of the Royal Society Interface, 12(104):20141088, 2015.

[39] Mark Jankauski, TL Daniel, and IY Shen. Asymmetries in wing inertial and aerodynamic torques contribute to steering in flying insects. Bioinspiration & biomimetics, 12(4):046001, 2017.

[40] Sanjay P Sane and Michael H Dickinson. The aerodynamic effects of wing rotation and a revised quasi-steady model of flapping flight. Journal of experimental biology, 205(8):1087–1096, 2002.

[41] Toshiyuki Nakata, Hao Liu, and Richard J Bomphrey. A cfd-informed quasi-steady model of flapping-wing aerodynamics. Journal of fluid mechanics, 783:323–343, 2015.

[42] Q Wang, JFL Goosen, and F van Keulen. An efficient fluid–structure interaction model for optimizing twistable flapping wings. Journal of Fluids and Structures, 73:82–99, 2017.

